# Litmus-Body: a Molecularly Targeted Sensor for Cell-Surface pH Measurements

**DOI:** 10.1101/800003

**Authors:** Marc C. Goudge, Joe Chin-Hun Kuo, Ann E. Metzloff, Ling-Ting Huang, Marshall J. Colville, Warren R. Zipfel, Matthew J. Paszek

## Abstract

Precise pH measurements in the immediate environment of receptors is essential for elucidating the mechanisms through which local pH changes associated with diseased phenotypes manifest into aberrant receptor function. However, current pH sensors lack the molecular specificity required to make these measurements. Herein we present the Litmus-body, our recombinant protein-based pH sensor, which through fusion to an anti-mouse IgG nanobody is capable of molecular targeting to specific proteins on the cell surface. By normalizing a pH-dependent green fluorescent protein to a long-Stokes shift red fluorophore or fluorescent protein, we readily report pH independent of sensor concentration using a single 488-nm excitation. Our Litmus-body showed excellent responsiveness in solution, with a greater than 50-fold change across the physiological regime of pH. The sensor was further validated for use on live cells, shown to be specific to the protein of interest, and was able to successfully recapitulate the numerous pH changes along the endocytic pathway.

---

Acidification of the extracellular microenvironment is a hallmark of cancer progression^1,2^. In response to an increased anabolic demand associated with uncontrolled proliferation, cancer cells upregulate glycolysis and consequently overproduce protons intracellularly^3^. This excess of protons is then expelled to maintain cellular homeostasis, lowering the pH of the extracellular space. The consequences of extracellular acidosis are diverse and profound: for example, mutant receptors may become permanently activated in a low pH environment^4^, and acid-adapted cells show a proclivity towards a more aggressive phenotype^5^. While these effects have been studied in the context of the bulk extracellular pH^6,7^, work towards elucidating the precise relationship between localized acidity and aberrant cellular function has been limited by the ability to measure pH at precise locations.

Notably, this localized acidity is further exaggerated at the cancer cell surface^8^. It is suggested that cancer-associated proton secretion creates a concentration gradient of protons that is highest at the cell membrane^9,10^. In addition, negatively charged residues in the glycocalyx, a carbohydrate-enriched cell coat near the pericellular surface, have also been predicted to accumulate protons that lower the local pH^11^. In concert with cancer-associated acidosis, the content of anionic moieties in the glycocalyx has been shown to increase^12^, which could act synergistically with proton secretion to concentrate protons in the vicinity of cell surface receptors. These factors may lead to heterogeneity in the local pH that receptors experience, significantly impacting their functionality. Although cell surface pH can be determined by recently developed pH sensitive fluorescent dyes conjugated to pH-low insertion peptides^8^ and cell penetrating peptides^13^, there remains an unmet need in the development of molecularly targeted sensors capable of reporting on the immediate environment of receptors.

Molecular targeting can be achieved by the diverse repertoire of antibody specificity^14^. Monoclonal primary antibodies that target specific cell surface receptors are clinically important for cancer therapies, such as Cetuximab for the cancer types that overexpress epidermal growth factor receptor (EGFR)^15^. Primary antibodies are conventionally detected by a secondary antibody, although its relatively large size (∼150 kDa) can be disadvantageous in sample penetration and signal localization^16^. Recently, nanobodies, or single-domain antibodies, have emerged as attractive alternatives for specific binding and localisation to primary antibodies^17^. Specifically, Pleiner et al. have successfully expressed and validated a library for targeting rabbit IgG and all mouse IgG subclasses^18^. Their small size (∼13 kDa), ease of labelling and consistent behaviour in recombinant protein fusions make nanobodies well-suited for fluorescence imaging^18–20^.

Fluorescent protein-based biosensors are versatile tools that can be fused to nanobodies for specific targeting^21^. Through extensive protein engineering, derivatives of the green fluorescent protein (GFP) from *Aequorea victoria* have been selected for pH dependent fluorescence^22–25^. In particular, pHluorins have been widely used as genetically-encoded sensors for tracking the pH of intracellular compartments^22,24^. Notably, the bright 488 nm excitable superecliptic pHluorin (SEP) variant is a pH sensor that shows a ∼ 50-fold signal change between pH 5.5 and 7.5, making it ideal for applications in physiological conditions. One major drawback is the inability to distinguish pH-dependent fluorescence changes from the local variations in its concentration. Without signal normalisation, studies using SEP as a pH sensor have remained qualitative^22^.

On the other hand, the ratiometric variants exhibit two pH-sensitive UV excitation peaks that can be normalized against each other^22^. This feature is an important advantage over the attributes of SEP as it allows for pH quantification independent of sensor concentration. However, the utility of pHluorin and its fluorescence enhanced variant, pHluorin2, suffers from exposing cells to phototoxic UV light and reporting a limited dynamic range of only ∼3-fold signal change in physiological conditions^22,26^.

Consequently, we aim to take advantage of the superior pH sensitivity of SEP for pH quantification by normalising its response against a second fluorophore that displays a large-Stokes shift (LSS). Adopting this strategy could potentially allow the pH response of SEP to be normalised by a single-wavelength co-excitation at 488 nm, which, in addition to allowing for signal quantification, avoids the photocytotoxicity conferred by UV light excitation. The use of SEP in conjunction with LSS red fluorophores has the added benefit of allowing extra fluorophores to be excited by 594 nm and 633 nm confocal laser lines for additional multicolour imaging.

In this work, we describe the Litmus-body, a tandem protein fusion that incorporates an IgG-specific nanobody and an SEP-based sensor that can normalise its pH response to LSS fluorophores with a single-wavelength excitation. We show here that, as a proof-of-principle, the Litmus-body can be successfully targeted to IgG antibodies and provide pH measurements when localised to components of interests on the cell surface, as well as following their transit through the endocytic pathway.

## EXPERIMENTAL SECTION

### Antibodies and reagents

The following antibodies were used for immunostaining: mouse anti-human Muc1 (CD227) monoclonal antibody (555925; BD Biosciences), goat anti-mouse Alexa Fluor 647 (A-21236; Thermo Fisher Scientific) and mouse anti-human EGFR antibody (225/Cetuximab, MA5-12880; Thermo Fisher Scientific). Doxycycline (sc-204734; Santa Cruz) was used for human cell culture induction, and IPTG (14213-261; IBI Scientific) was used for bacterial culture induction. Kanamycin sulfate (420311; MilliporeSigma) was used for bacterial culture selection. Hoescht 33342 (H1399; Thermo Fisher Scientific) was used for nuclear staining. Normal goat serum (NGS; S1000; Vector Laboratories) was used as a blocking agent. The following buffers were prepared: 2.5X Ni-NTA binding buffer (375 mM NaCl, 125 mM K_2_HPO_4_, 25 mM Tris pH 8.5, 25 mM imidazole), Ni-NTA wash buffer (300 mM NaCl, 50 mM K_2_HPO_4_, 20 mM imidazole), Ni-NTA equilibration buffer (300 mM NaCl, 50 mM K_2_HPO_4_, 10 mM imidazole), Ni-NTA elution buffer (300 mM NaCl, 50 mM K2HPO4, 250 mM imidazole), and maleimide labelling buffer (MLB; 100 mM K_2_HPO_4_, 150 mM NaCl, 1 mM EDTA, 250 mM sucrose). Unless otherwise stated, all chemicals were purchased from MilliporeSigma, cell culture reagents were purchased from Thermo Fisher Scientific and enzymes for molecular cloning were purchased from New England Biolabs (NEB).

### Construct generation

A dsDNA oligo encoding a cysteine-free SEP (cfSEP) engineered with an additional C-terminal surface cysteine (IDT), using NEB HiFi Assembly, was inserted into a BamHI-HF and NcoI-HF linearized pTP1112 vector (generously provided by Dirk Görlich: Addgene plasmid #104158)^18^. pTP1112 encodes an N-terminal His_14_-*bd*NEDD8 tagged anti-mouse IgG1 Fc specific TP1107 nanobody with an ectopic C-terminal cysteine. This terminal cysteine was removed and replaced with a Gly-Gly-Gly-Gly-Ser flexible linker, to ensure that downstream cysteine-maleimide labelling only occurred at the cfSEP C-terminal cysteine and to improve folding of the fusion protein. These steps generated a His_14_-*bd*NEDD8-TP1107-cfSEP construct.

mCyRFP1 was extracted from a mMaroon-mCyRFP1-pET11(a) construct (unpublished work) using the Q5 Hot Start High-Fidelity Master Mix (NEB) with 5’-ATGAACTGTACAAAGGAGGAGGCGGTAGCATGGTTAGTAAAGGCGAAGAAC-3’ (forward) and 5’-CCAAGCTCAGCTAAAGCTTATTTATACAGTTCATCCATGC-3’ (reverse). The His_14_-*bd*NEDD8-TP1107-cfSEP construct was linearized with Gibson Assembly compatible ends (forward: 5’-TAAGCTTTAGCTGAGCTTGGAC-3’, and reverse 5’-CATGCTACCGCCTCCTCCTTTGTACAGTTCATCCATG-3’ (note: the engineered cfSEP C-terminal cysteine was removed and replaced with a Gly-Gly-Gly-Gly-Ser flexible linker). Afterwards, the linear fragments were combined via NEB HiFi Assembly to form a His_14_-*bd*NEDD8-TP1107-cfSEP-mCyRFP1 construct.

Single fluorescent proteins were generated via Q5 site-directed deletion of the His_14_-*bd*NEDD8-TP1107-cfSEP-mCyRFP1 construct. *bd*NEDD8, TP1107 nanobody, and the unneeded fluorescent protein were deleted using the following primer pairs: 5’-ATGGTTAGTAAAGGCGAAGAACTGATT-3’ (forward) with 5’-TGATCCGCCGGTATGGTGATGACT-3’ (reverse) to isolate a His_14_-mCyRFP construct, and 5’-ATGGTGAGTAAAGGAGAAGAACTTTTCACTGG-3’ (forward) with 5’-TGATCCGCCGGTATGGTGATGACT-3’ (reverse) to isolate a His_14_-cfSEP construct.

His_14_-SEP was made by a two-fold site-directed mutagenesis of the His_14_-cfSEP construct in order to generate linear fragments compatible for NEB HiFi Assembly. The S48C mutation was generated using 5’-CTTACCCTTAAATTTATTTGCACTACTGGAAAACTACC-3’ (forward) and 5’-TGATCTGGGTATCTTGAAAAGCATTGAACACCATAAGT-3’, and the M70C mutation was generated with 5’-ACTTATGGTGTTCAATGCTTTTCAAGATACCCAGATCA-3’ (forward) and 5’-GGTAGTTTTCCAGTAGTGCAAATAAATTTAAGGGTAAG-3’ (reverse). The two linear strands were subsequently joined.

### Cell lines and cultures

All cells were maintained with 100 U/mL Penicillin/Streptomycin at 37 °C, 90% relative humidity, and 5% CO2. A previously described MCF10A cell line that stably expressed a doxycycline-inducible rtTA NeoR Mucin-1 (Muc1) deficient of cytoplasmic tail (dCT) was cultured in DMEM/F12 media supplemented with 5% horse serum, 20 ng/mL EGF (Peprotech), 10 mg/mL insulin, 500 ng/mL hydrocortisone, and 100 ng/mL cholera toxin^27^. A431 cells (ATCC) were cultured in DMEM supplemented with 10% FBS, 1 mM pyruvate, and 1X Glutamax.

### Recombinant protein production and purification

All recombinant proteins were expressed in chemically competent NiCo21 (DE3) *E. coli* (NEB). 5 mL precultures (LB containing 50 µg/ml kanamycin) were grown overnight at 37 °C, 220 rpm and then diluted with fresh medium to 0.2 -1 L, in baffled flasks at no more than a 1:10 media:container volume ratio, and allowed growth to reach OD600 of just below 0.6. The cultures were induced by 0.5 mM IPTG overnight at 24 °C, harvested, resuspended in B-PER (Thermo Fisher Scientific) and vortexed for cell lysis. The lysates were cleared by centrifugation at 10,000 g for 20 min at 4 °C.

His_14_-tagged recombinant proteins were purified using immobilized metal affinity chromatography. Supernatant diluted into 1X Ni-NTA binding buffer was bound to equilibrated Ni-NTA resin for 20 min at 4 °C, with end-over-end mixing. The resin was added to a spin column, washed thoroughly and incubated with the Ni-NTA elution buffer for 20 min at 4 °C, with end-over-end mixing. Recombinant proteins were then eluted and buffer exchanged (fluorescent proteins to pH 7.4 PBS, and *bd*NEDP1 to 50 mM Tris-HCl, 300 mM NaCl, 250 mM sucrose, 10 mM DTT, pH 7.5) using Zeba 7k MWCO desalting columns or by overnight dialysis with 10k MWCO Snakeskin dialysis tubing. Eluted proteins were then sterile filtered and snap-frozen for long-term storage at -80 °C. Nanobody-containing constructs were mixed with 0.1% w/v sodium azide prior to snap-freezing.

### NEDD8-removal

*bd*NEDP1 protease expressed from the pDG02583 construct (a gift from Dirk Görlich; Addgene plasmid # 104129) was used to remove *bd*NEDD8^28^. Ni-NTA purified His_14_-*bd*NEDD8-tagged proteins were incubated with >500 nM of the protease for 2 h on ice. The protease-protein mixture was incubated with equilibrated Ni-NTA resin for 20 min and spun at 700 g to elute the cleaved protein while leaving the uncleaved protein and the protease bound to the column. The cleaved protein was then buffer exchanged using a Zeba 7k MWCO desalting column to Ph 7.4 PBS, prior to filtration and snap-freezing.

### ATTO490LS-maleimide labelling

Cysteine-maleimide labelling was performed as previously described and all steps were kept on ice to protect internal cysteines from the labelling reaction^20^. Briefly, the engineered cfSEP C-terminal cysteine was reduced by 15 mM TCEP for 10 mins. TCEP was removed by buffer exchange to degassed MLB using a Zeba 7k MWCO desalting column (Thermo Fisher Scientific). ATTO490LS-maleimide (ATTO-TEC Gmbh) was added to the reduced TP1107-cfSEP at a 6:5 molar ratio and the reaction-mixture was brought to pH 7.5 with K_2_HPO_4_. The mixture was stirred on ice under nitrogen for 1.5 h. Excess ATTO490LS was removed through buffer exchange to MLB.

### SYPRO Ruby protein gel staining

Proteins were diluted with 4x NuPAGE LDS Sample Buffer (Thermo Fisher Scientific) and heated at 70°C for 10 min. Proteins were subsequently separated by SDS-PAGE on a 4-12% NuPAGE BisTris pre-cast gradient gel (Thermo Fisher Scientific) at 200V for 35 min. Gels were fixed, stained with SYPRO Ruby Protein Gel Stain (Thermo Fisher Scientific), and washed according to manufacturer’s specification. Washed gels were imaged on a ChemiDoc MP Imaging System (Bio-rad).

### Photophysical properties

Quantum yield was determined at 490 nm in PBS, pH 7.4, by acquiring integrated fluorescence (500 nm – 800 nm) in conjunction with absorbance values in a dilution series from A_490_ ∼ 0.1, to minimize inner filtering effect, using fluorescein as a standard (Quantum yield = 0.925 at 0.1 M NaOH)^29^. Molar extinction coefficient was determined by measuring mature chromophore concentration under NaOH denaturing conditions. Absorbance at 450 nm was measured immediately after mixing proteins with equal volume of 2 M NaOH^30,31^. This assumed alkali-denatured chromophore exhibited extinction coefficient 44,000 M^-1^ cm^-1^ at 450 nm absorbance. Total protein concentration was determined at 280 nm absorbance. All absorbance was measured on a Cary 300 UV-VIS spectrometer (Agilent Technologies Inc.) and fluorescence spectra were recorded by a QM4 fluorimeter (Horiba Instruments/Photon Technology International). Brightness was calculated as the product between quantum yield and extinction coefficient.

Fluorescence lifetimes were measured by time-correlated single photon counting (TCSPC). Sample solutions were excited by a 445 nm picosecond diode laser (BDL-445-SMC, Becker & Hickl GmbH) pulsed at a 20 MHz repetition rate. Fluorescence decay curves were collected from samples in 1 cm pathlength quartz cuvettes at 90 degrees to the 445 nm excitation using a R3809U-50 microchannel plate photomultiplier tube with a 25 ps transit time spread (Hamamatsu, Bridgewater, NJ). Excitation intensity was attenuated using a Glan-Thompson polarizer to keep the phospholuminescence detection rate less than 0.2% of the repetition rate to avoid photon pile-up. Data was acquired using a TCSPC module (SPC-830, Becker & Hickl GmbH) and fit to a bi-exponential decay using the SPCImage software (Becker & Hickl GmbH).

### Solution pH response

Universal buffer solutions were prepared according to the Carmody buffer series^32^. This involved mixing a master acid buffer (0.2 M Boric Acid, 0.05 M Citric Acid) and a master base buffer (0.1 M NaHPO_4_) at previously determined ratios to achieve approximate pH of interest. Fluorescent constructs were diluted to 100 nM in these buffers, and the pH of the buffer-protein solution was recorded using an Orion™ PerpHecT™ ROSS™ Combination pH Micro Electrode (Thermo Fisher Scientific). The buffer-protein solutions had their emission spectra measured using a Tecan 1000M Infinite plate reader at 490 nm excitation. Three sets of triplicates were recorded for each bufferprotein solution. pKa and Hill coefficient were determined by least square non-linear fitting the normalized data to the five-parameter logistic Richards equation on GraphPad Prism (California USA). Fold change was calculated as the intensity_pH7.5_/intensity_pH5.5_, where the values at specific pH were determined through linear interpolation.

### Cell binding assay

Doxycycline-induced MCF10A Muc1dCT cells were fixed with 4% paraformaldehyde for 10 min at room temperature and blocked in PBS containing 5% NGS for 1 h at room temperature. All subsequent dilutions were performed in PBS containing 5% NGS. As a positive control, cells were incubated with the primary anti-Muc1 antibody at 1:1,000 dilution overnight at 4 °C and detected using the secondary Alexa Fluor 647-labelled goat antimouse IgG at 1:1,000 dilution for 1 h at room temperature. To test Litmus-body as a secondary reagent, cells were incubated with the primary anti-Muc1 antibody at a 1:1,000 dilution overnight at 4 °C and then incubated with 30 nM of the Litmus-body for 1 h at room temperature. For one-step immunostaining, Litmus-body was pre-incubated with the anti-Muc1 antibody at equimolar ratio overnight at 4 °C. 30 nM of the pre-incubated IgG-Litmus-body complex was then applied to cells for 1 h at room temperature. For negative controls, cells were labelled with 30 nM Litmus-body for 1 h at room temperature in the absence of a primary IgG antibody. All samples were labelled with Hoechst at 1 µg/mL for 15 min. Cells were then imaged on a Zeiss 800 LSM microscope using a 20X air objective (NA 0.8).

### Muc1 live cell imaging

Mouse anti-human Muc1 monoclonal antibody and Litmus-body were mixed at equimolar ratio, 4 °C, overnight, to form an IgG-Litmus-body complex. Doxycycline-induced MCF10A Muc1dCT cells were incubated at 37 °C for 30 min with PBS containing 5% NGS and 0.1% sodium azide to inhibit endocytosis. Cells were then incubated further with 33 nM of the IgG-Litmus-body complex at 37 °C for 30 min, washed and imaged in PBS containing 5% NGS and 0.1% sodium azide with pH adjusted to 6, 7.03 and 7.95 by an Orion™ PerpHecT™ ROSS™ Combination pH Micro Electrode (Thermo Fisher Scientific). Spectral imaging was performed using lambda mode, at 488 nm excitation and 9 nm spectral resolution, on a Zeiss LSM880 inverted confocal microscope with a 40X water objective (NA 1.1).

### Quantification of lambda mode images

Thresholding was applied to the 22 channel, 16-bit Lambda-stacked images based on the pixel intensity values of the cfSEP and mCyRFP1 emission channels (cfSEP channel wavelength: 513 nm, mCyRFP1 channel wavelength: 593 nm). Pixels below the threshold (750 AU) in either of the two channels were ignored for the calculation. Mean of the non-thresholded pixels was subsequently used for calculating the average spectra of the selected image subsets, as well as for normalizing the displayed stacks.

### Endocytosis studies

Mouse anti-human EGFR monoclonal antibody (225/Cetuximab) and Litmus-body were mixed at equimolar ratio, 4 °C, overnight, to form an IgG-Litmus-body complex. To generate a calibration curve, the IgGLitmus-body complex was diluted to a final concentration of 200 nM in PBS containing 5% NGS and adjusted to final pH values of 4, 5.04, 5.49, 6.09, 6.56, 7.01, 7.49, 7.91 using an Orion™ PerpHecT™ ROSS™ Combination pH Micro Electrode (Thermo Fisher Scientific). Least square non-linear fitting was performed using the four-parameter logisticmodel on GraphPad Prism (California USA). IgG-Litmus-body fluorescence in each pH buffer was then acquired on a Zeiss LSM800 inverted confocal microscope using a 63 X water objective (NA 1.2). The microscope was configured to simultaneously scan for cfSEP and mCyRFP1 signal with 488 nm excitation. To minimize cross-talk, the emission bands were set to collect between 500 – 535 nm for cfSEP and 575 – 700 nm for mCyRFP1.

For intracellular trafficking experiments, A431 plated overnight at 10,000 cells/cm^2^ were chilled on ice for 1 h to slow down endocytosis, washed and incubated with 67 nM of the IgG-Litmus-body complex in PBS containing 5% NGS at 37 °C for 30 min. Cells were washed and allowed to undergo endocytosis for 1 h at 37 °C. They were further washed and placed in a pH 7.5 PBS buffer containing 5% NGS for imaging. Cells were then placed in a pH 5.2 PBS buffer containing 5% NGS and imaged. Cell imaging was done on a Zeiss LSM800 inverted confocal microscope using a 63 X water objective (NA 1.2) and Litmus-body imaging used the same configuration for generating the calibration curve above.

## RESULTS AND DISCUSSION

### Litmus-body and expression

We set out to create Litmus-body, a nanobody-sensor fusion, that can be specifically targeted to mouse IgG antibodies to report on the local pH of cell surface components. Sensor fusion proteins were produced in *E. coli* for ease of culture and scaling. We took advantage of a highly soluble protease-cleavable tag previously reported by Frey and Görlich (*bd*NEDD8) to optimise expression of the nanobody component in our sensor^28^. Removable by its associated protease *bd*NEDP1, the *bd*NEDD8 tag has been shown to improve the cytoplasmic yield of nanobodies in *E. coli* (Supplemental Figure 1). To generate an IgG specific Litmus-body, we fused fluorescent sensor proteins to *bd*NEDD8-TP1107, an anti-mouse IgG Fc fragment nanobody, that allowed the pH sensor to be applied in a manner analogous to secondary antibodies upon cleavage of NEDD8 (Figure 1A).

**Figure 1.**
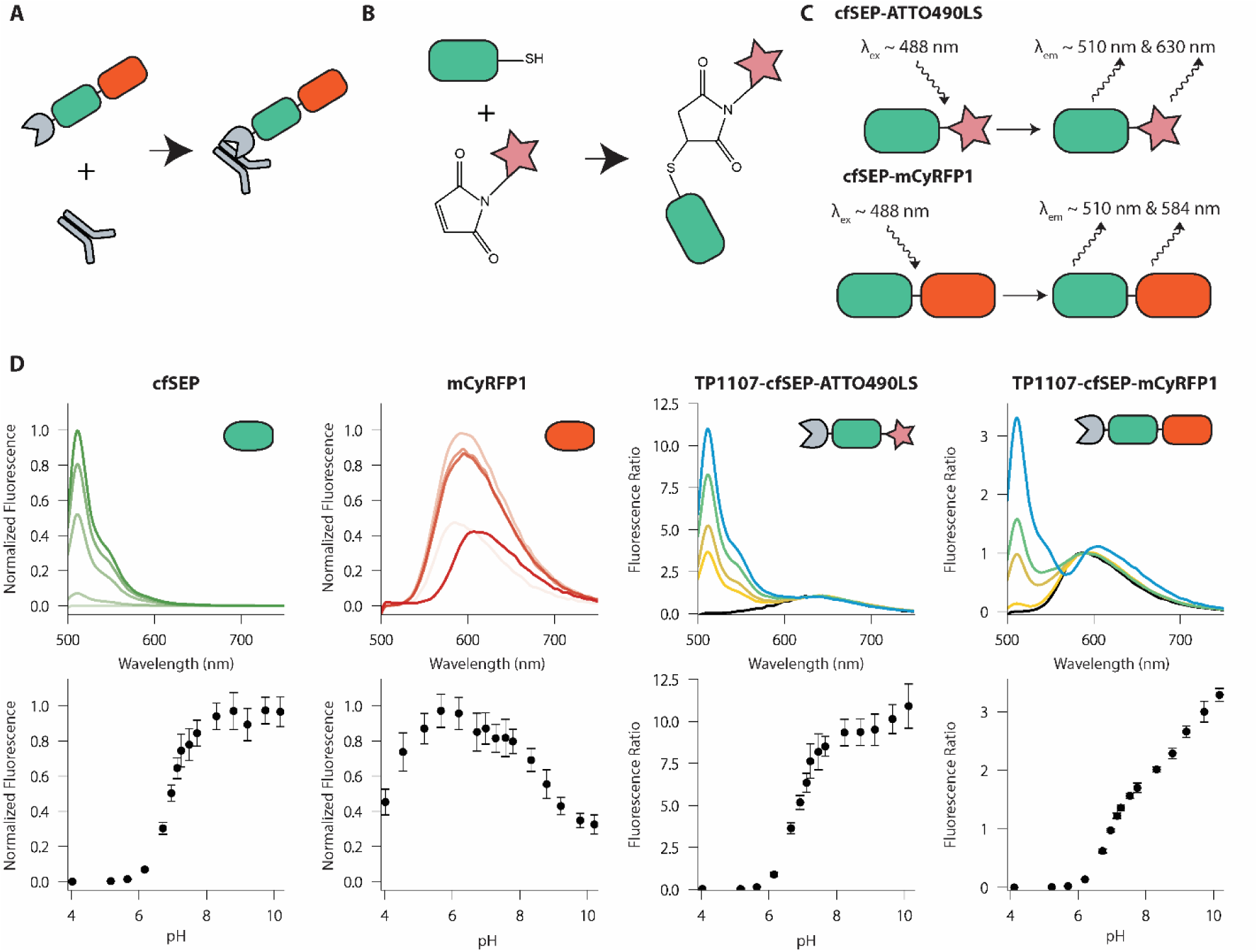
Litmus-body design and solution properties. A) Overview of the Litmus-body. A pH-sensitive GFP derivative (green) and red fluorophore (red) are fused to an anti-IgG nanobody (grey), allowing for targeting of primary antibodies. B) Reaction scheme for the conjugation of cfSEP to a maleimide-bound fluorophore. C) Scheme of the dual emission nature of the Litmus-body, using both ATTO490LS (top) and mCyRFP1 (bottom) as the long Stokes-shift red fluorescence emitter for ratiometric normalization of green sensor emission. red fluorophore. D) Fluorescence spectra and pH responsiveness of TP1107-cfSEP-mCyRFP1, TP1107-cfSEP-ATTO490LS, and their constituent fluorescent proteins. The fluorescence ratio for TP1107-cfSEP-mCyRFP1 was calculated as (I510/I590), while the fluorescence ratio for TP1107-cfSEP-ATTO490LS was calculated as (I510/I630). Error bars represent standard deviation of three independent experiments, each with triplicate.

### Design and validation of a cysteine-free SEP engineered for maleimide chemistry

To explore the potential use of a synthetic dye for pH signal normalisation, we created a cysteine-free SEP (cfSEP) engineered with an ectopic C-terminal cysteine for site-specific maleimide-dye labelling (Figure 1B). Unlike conventional non-selective modifications, such as via N-hydroxysuccinimide ester, maleimide reaction on C-terminal cysteines has been demonstrated to be an effective protein labelling strategy that produces homogenous protein-dye-ratio conjugates, reduces batch-to-batch variation and does not alter binding properties of the modified protein^20,33^.

We introduced C48S/C70M mutations to remove the native cysteines in SEP and generated cfSEP. This was to avoid off-target cysteine-maleimide reactions and to ensure that only the engineered C-terminal cysteine would be available for dye-conjugation. C48S/C70M mutations have previously been validated in GFP derivatives and shown to maintain fluorescence performance in EGFP, a pH sensitive protein with similar characteristics to SEP^34^. cfSEP with the engineered C-terminal cysteine was tested for its pH response. It retained similar idea pKa and responsiveness to SEP in the physiological range, while also displaying consistent photophysical properties to SEP (Table 1, Supplemental Table 1, Supplemental Figure 2A). These results pointed to the suitability of cfSEP as a substitute for SEP.

**Table 1.**
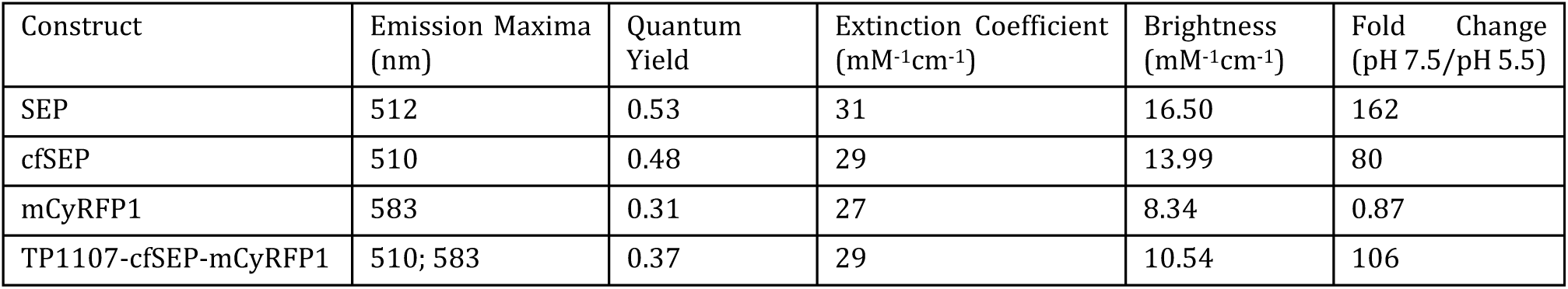
Photophysical properties and pH response characteristics of fluorescent proteins and constructs.

| Construct | Emission Maxima (nm) | Quantum Yield | Extinction Coefficient ($\text{mM}^{-1}\text{cm}^{-1}$) | Brightness ($\text{mM}^{-1}\text{cm}^{-1}$) | Fold Change (pH 7.5/pH 5.5) |
| --- | --- | --- | --- | --- | --- |
| SEP | 512 | 0.53 | 31 | 16.50 | 162 |
| cfSEP | 510 | 0.48 | 29 | 13.99 | 80 |
| mCyRFP1 | 583 | 0.31 | 27 | 8.34 | 0.87 |
| TP1107-cfSEP-mCyRFP1 | 510; 583 | 0.37 | 29 | 10.54 | 106 |

### Expression and pH response of TP1107-cfSEPATTO490LS

ATTO490LS-maleimide was conjugated to TP1107-cfSEP on the engineered C-terminal cysteine to generate a TP1107-cfSEP-ATTO490LS construct (Figure 1B). ATTO490LS is a long-stoke shift synthetic dye with peak excitation at 496 nm. These properties make ATTO490LS an attractive normalisation partner for cfSEP as both fluorophores can be simultaneously excited and individually resolved (Figure 1C). We recorded the fluorescence response to solution pH of the TP1107-cfSEP-ATTO490LS construct (Figure 1D). The response of the conjugate at 510 nm emission (Supplemental Figure 2C) showed a similar pKa and responsiveness to the unconjugated cfSEP (Supplemental Table 1), indicating that fusion and labelling of cfSEP did not negatively impact functionality. pH response of TP1107-cfSEP-ATTO490LS was determined by normalising cfSEP signal against ATTO490LS (Figure 1D). The construct readily detected pH changes in the physiological regime, exhibiting >50-fold signal enhancement from pH 5.5 to 7.5. Interestingly, we noted that the 630 nm emission in the construct increased at a rate greater than both the cfSEP and ATTO490LS individually (Supplemental Figure 2B, 2C), potentially indicating the occurrence of fluorescence resonance energy transfer (FRET) between cfSEP and ATTO490LS. We subsequently found that the fluorescence lifetime of the combined construct (∼1.5 ns) was lower than that of cfSEP (∼2 ns), further pointing to the occurrence of FRET.

### Expression and pH response of TP1107-cfSEP-mCyRFP1

To avoid the complication of FRET in our construct, we looked to monomeric cyan-excitable red fluorescent protein (mCyRFP1) instead as a normalisation partner for cfSEP. mCyRFP1 is a long-stokes shift TagRFP derivative with broad-excitation around 500 nm^30^. Like ATTO490LS, mCyRFP1 can be co-excited with cfSEP while displaying an easily separable emission. The C-terminal cysteine on cfSEP was removed and replace by a flexible GGGGS peptide linker to fuse to mCyRFP1. We recorded the fluorescence response to solution pH of the construct. The 510 nm emission of the construct showed a pKa and responsiveness similar to the unconjugated cfSEP (Supplemental Figure 2D, Supplemental Table 1), suggesting that fusion to a red fluorophore did not impact cfSEP performance. mCyRFP1 showed a distinctive and consistent pH response, which was retained in the 590 nm emission of the construct (Figure 1D, Supplemental Figure 2D).

The pH response of TP1107-cfSEP-mCyRFP1 was determined by normalising cfSEP signal against mCyRFP1’s contribution (Figure 1D). TP1107-cfSEP-mCyRFP1 performed similarly to TP1107-cfSEP-ATTO490LS, also exhibiting >50-fold signal increase from pH 5.5 to 7.5. Moreover, the mCyRFP1 fused construct benefited from the additional pH responsiveness of mCyRFP1 above pH 8. This supplemented the overall responsiveness of the sensor upon the saturation of the cfSEP signal and extended the range of pH sensitivity in more basic environments (Figure 1D). We thus moved forward with the TP1107-cfSEP-mCyRFP1 construct for cellular testing and hereinafter referred to it as the Litmus-body.

### Molecular targeting and response to environmental pH on live cells

Litmus-body was targeted to specific cancer cell surface components by its ability to piggyback on IgG antibodies. Mucin-1 (Muc1) was selected as an epitope of interest given its key role in forming the cancer cell glycocalyx^27,35,36^. Fixed Muc1-overexpressing cells stained with a primary anti-Muc1 IgG antibody could be similarly detected either by a secondary antibody or by using the IgG-specific Litmus-body as a secondary reagent (Figure 2A). We also allowed the Litmus-body to react to the primary anti-Muc1 IgG antibody at an equimolar ratio before cell application to optimize incubation time for live cell applications. The IgG-Litmus-body complex exhibited a similarly high specific binding to the surface of Muc1-overexpressing cells (Figure 2B). These results verified that the specificity of TP1107 binding to IgG antibodies was unaffected in the sensor construct and that Litmus-body could be used as a simple, one-step targeting reagent by pre-complexing with an IgG antibody.

**Figure 2.**
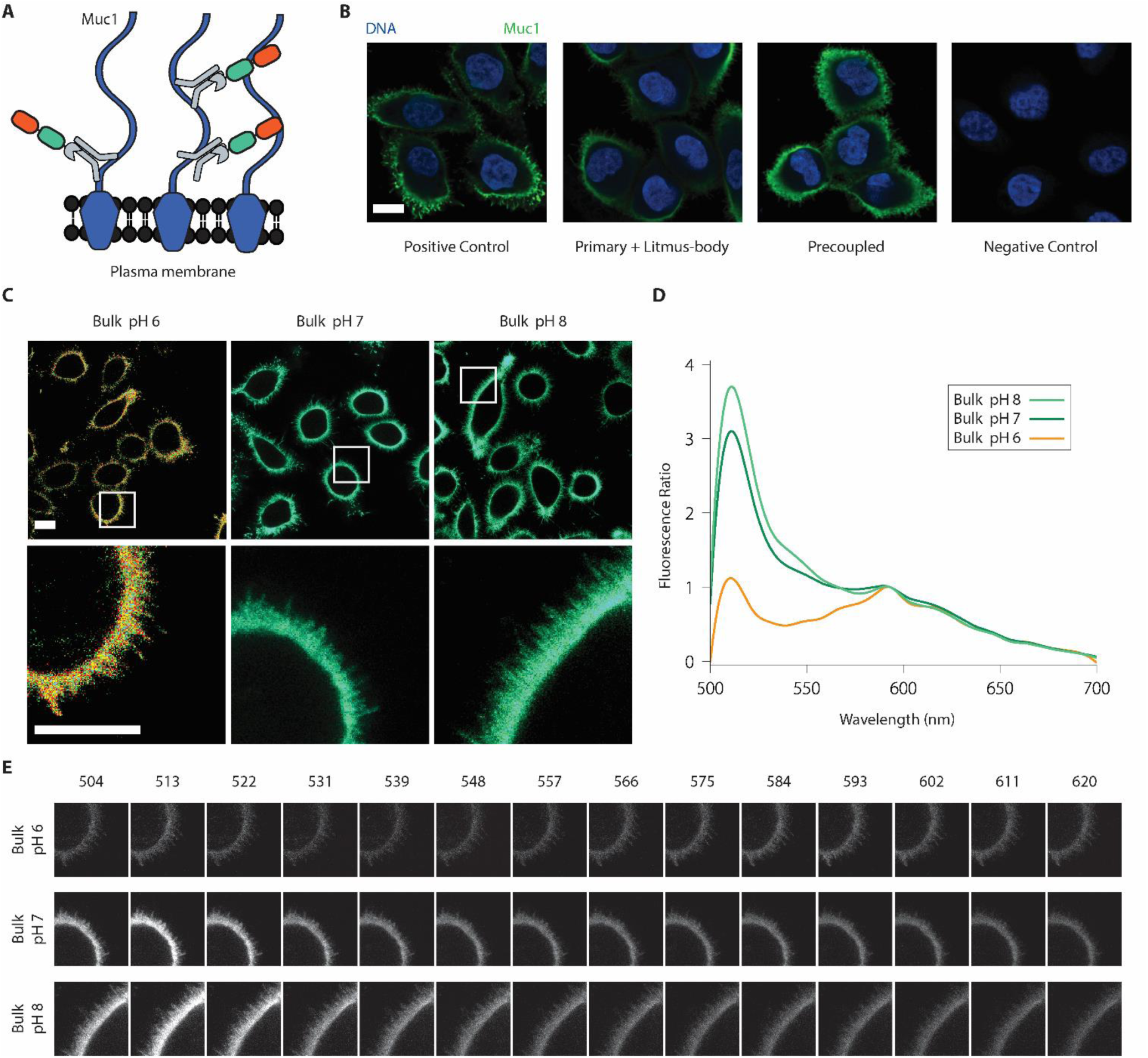
Litmus-bodies bind specifically to their epitope of interest and respond to bulk pH change on live cells. A) Scheme depicting the nature of binding between Muc1, and precoupled anti-Muc1-Litmus-Body on Muc1 overexpressing MCF10As. Numerous anti-Muc1 binding epitopes are present along the cell surface, allowing for dense labelling. B) Representative immunofluorescence images from three independent experiments depicting the specificity of the Litmus-body as compared against conventional secondary antibody staining (Blue: DNA, and Green: 488 nm excitation) on paraformaldehyde fixed Muc1 overexpressing MCF10A cells. Positive control depicts treatment with an anti-Muc1 primary and Alexa Fluor 488 anti-IgG secondary. Cells were treated with an anti-Muc1 primary and the Litmus-body was used as a secondary reagent (Primary + Litmus-body). To test for a simple, one-step staining procedure, anti-Muc1 antibody was pre-reacted with the Litmus-body to form an anti-Muc1-Litmus-Body complex before applying to cells (Precoupled). C) Representative live cell images from three independent experiments of pre-coupled anti-Muc1-Litmus-body binding to Muc1 overexpressing MCF10A cells treated with 0.1% (w/v) sodium azide in various bulk pHs. The bottom row depicts select regions of interest (white boxes). D) Fluorescence spectra normalized against the mCyRFP1 emission (calculated as (I/I_593_)), from the region of interest presented in C). E) Spectral stacks of the three regions of interests depicted in C), with the wavelength listed in nm. Panels for each sample were normalized against their mean intensity at 593 nm. Scale bars: 10 μm.

The IgG-Litmus-body complex was targeted to the surface of live Muc1-overexpressing cells and tested for its response to environmental pH. We treated these cells with sodium azide to inhibit endocytosis and minimize the internalization of the pH sensor. cfSEP and mCyRFP1 were simultaneously excited with a 488 nm laser and spectrally imaged. Emission peaks were observed in normalized spectra around the expected 510 nm and 583 nm for cfSEP and mCyRFP1, respectively. Increasing bulk solution pH from 6 to 8 brought about a concurrent signal increase of cfSEP relative to mCyRFP1 on the cell surface (Figure 2C, 2D, 2E). These results suggested that the IgG specific Litmus-body could be readily targeted to the cell surface and may act as a suitable agent to report local pH perturbations on the surface of live cells.

### pH imaging of cell surface receptor following endocytosis

We further explored the cellular applications of the Litmus-body by following its intracellular trafficking after binding to epidermal growth factor receptor (EGFR), an overexpressed drug target on multiple cancer types^37^. The endocytic pathway provides an ideal environment for validating our Litmus-body, given its diverse range of pH values that are well-characterized in cellular compartments along the pathway^38^. Confocal imaging experiments were configured to simultaneously excite and collect the emission of both cfSEP and mCyRFP1 to avoid excessive photobleaching. As above, for simple one-step targeting, Litmus-body was first reacted to a monoclonal anti-EGFR IgG antibody. We selected Cetuximab/C225 anti-EGFR antibody for this purpose due to its clinical importance for treating multiple cancer types including skin, colorectal, head and neck^39,40^. Cetuximab blocks ligand binding to EGFR and induces receptor mediated endocytosis^41^. After reacting the Litmus-body to Cetuximab, a calibration curve was obtained on the confocal by curve fitting of cfSEP/mCyRFP1 fluorescence ratio in solution (Figure 3A).

**Figure 3.**
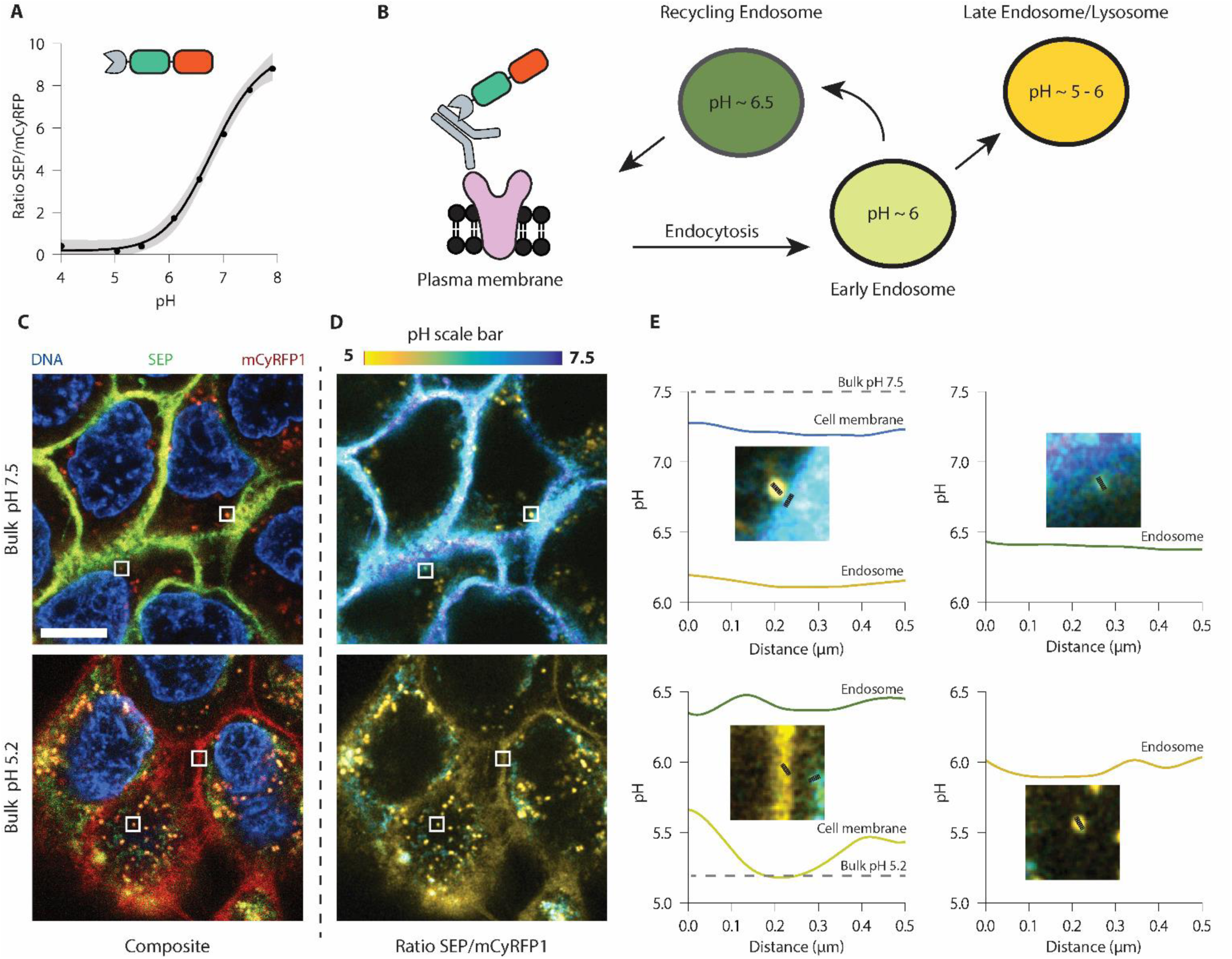
Litmus-bodies readily detect pH changes along the endocytic pathway. A) In-solution calibration of the Litmus-body based on (I_SEP_/I_mCyRFP1_) at various bulk pH. Points were fitted to a four parameter logistic function. Grey-shaded regions represented a 95% CI. B) Scheme depicting the nature of binding between epidermal growth factor receptor (EGFR), and pre-coupled anti-EGFR-Litmus-Body on EGFR overexpressing A431s. Upon binding to EGFR, the IgG-Litmus-body complex is endocytosed, allowing it to monitor changes in pH along the pathway. C) Representative live cell fluorescence images of pre-coupled anti-EGFR and Litmus-body binding to EGFR overexpressing A431s and being endocytosed. Top: bulk solution pH of 7.5. Bottom: Bulk solution of 5.2. D) Ratiometric (I_SEP_/I_mCyRFP1_) variants of the images presented in Figure C). Using the calibration curve from A), ratios were converted to pH values. E) Line traces (black lines) were taken from regions of interest in C/D) (white boxes, inline in E)). Scale bar: 10 μm.

The Cetuximab-Litmus-body complex was applied to A431 epidermoid cancer cells, which overexpressed EGFR by gene amplification^42^, and allowed to be internalized by receptor-mediated endocytosis at pH 7.5 (Figure 3B). We then switched the bulk solution to pH ∼5 to quench cfSEP signal on the cell surface. cfSEP and mCyRFP1 signals were observed in the composite images (Left panels, Figure 3C) and were processed to represent their fluorescence ratio (Figure 3D). These ratios were converted to pH values based on the calibration curve (Figure 3A & 3D). Interestingly when the bulk solution was adjusted to pH 7.5, regions on the membrane were observed at a slightly lower pH of ∼7.2 (Top panel, Figure 3D). This is consistent with previous reports suggesting that cancer cell surfaces have a lower pH than the bulk extracellular environment^8^.

In contrast, Litmus-body sequestered inside endocytic vesicles reported pH ranges from ∼5 to 6.5 (Figure 3D). Note that these Litmus-body-loaded intracellular vesicles were similar in size to endosomes and reported the expected pH values for these compartments (Figure 3B). At a bulk solution of pH 5.2, the cell surface pH reported by Litmus-body closely matched the bulk solution pH (Bottom panel, Figure 3D) though its membrane signal became less defined, possibly due to its dissociation from EGFR at low pH^43^. In contrast, the juxtaposed intracellular compartment reported pH ∼6.5 against the low cell surface pH. Altogether, these results suggested that our Litmus-body could be a useful tool for reporting the microenvironmental pH that its molecular targets may experience.

## CONCLUSION

Cell surface acidification is a hallmark of aggressive diseases such as cancer^1,2^. Here, we presented the Litmus-body, an IgG-specific pH sensor in which we fused together an anti-mouse IgG TP1107 nanobody, a pH responsive cysteine-free super-ecliptic pHluorin (cfSEP) and a large-stoke shift monomeric cyan-excitable red fluorescent protein (mCyRFP1), and demonstrated its ability to quantitatively monitor the local pH surrounding cell surface targets. Co-excitation and separable emission of cfSEP and mCyRFP1 made it possible to normalize the pH response of the Litmus-body by a single-wavelength excitation. By engineering cfSEP for maleimide labelling, we also described a synthetic dye-conjugate variant with a large-stoke shift ATTO490LS that replaced mCyRFP1 for signal normalization. The dye-conjugate would benefit from the reduced size of a synthetic dye (ATTO490LS; < 1 kDa) compared to a fluorescent protein (mCyRFP1; 26.4 kDa), though we observed considerable fluorescence resonance energy transfer (FRET) between cfSEP and ATTO490LS. Further improvements on the use of a 488 nm-excitable large-stoke shift dye without spectral overlap with cfSEP would minimise the complications of FRET. This highlights the utility of the modular design of the Litmus-body. As new fluorophores are developed, components may be swapped out for the current state of the art.

The modular design of the Litmus-body would allow the nanobody domain to be swapped for other targeting variants. While we focused our proof-of-principle studies on monoclonal anti-mouse IgG antibodies as their derivatives play important roles in therapeutic antibodies in the clinic^44,45^, we envisage that the anti-mouse IgG nanobody domain can be easily replaced by anti-IgG nanobodies that are currently available from other species including rabbit^18^. Furthermore, while unexplored in our work, a diverse palette of nanobodies have been developed for commonly expressed molecular targets including EGFR^46^. Litmus-body variants that can be directly localized to targets of interest would remove the need for a primary antibody, and further benefit from the small size of the Litmus-body to maximize cell and tissue sample penetration and perfusion.

Preformed IgG-Litmus-body complexes derived from reacting IgG antibodies to the Litmus-body provided a simple, time-effective one-step targeting strategy for live cell applications. This approach benefited from the monomeric and monovalent nature of anti-IgG nanobodies, as they do not cross-link primary antibodies to form large multimeric complexes that impede IgG binding^18^. Using this strategy, we targeted the Litmus-body to two oncogenic cell surface proteins that are overexpressed in multiple cancer types: Muc1 and EGFR^37,47^, and demonstrated that the Litmus-body responded well to pH changes surrounding these proteins. Importantly, the Litmus-body reported a decreased pH surrounding EGFR on the cancer cell surface compared to the bulk solution. This observation is consistent with recent reports in the literature that suggest the pH on cancer cell surfaces is lower than the bulk microenvironment^8^. The ability to target cell surface proteins is a major advantage of the Litmus-body over current strategies in the literature that lack molecular specificity, such as using low pH insertion peptides^8^, and paves the way to elucidating the mechanisms that give rise to their aberrant function in the cancerous phenotype. The ability of the Litmus-body to piggyback on the vast diversity and high specificity of IgG antibodies may prove to be a powerful approach that opens up an almost infinite number of molecular targets.

Antibody trafficking into acidic intracellular compartments may be important for payload release in the design of antibody-drug conjugates (ADCs) used in cancer therapies^48^. In our proof-of-principle studies, Litmus-body reported on the pH surrounding the internalized Cetuximab, an EGFR specific antibody used in cancer treatment that is undergoing active research for ADC based therapies^49^. This intracellular tracking was enabled by the single-wavelength co-excitable and dual-emission nature of the Litmus-body that permitted the accurate co-localisation of cfSEP and mCyRFP1. Simultaneous excitation of fluorophores with a single-wavelength light can minimize the effect of sample movement, uneven sensor distribution and sample thickness variations^50^. It also reduced image acquisition time, phototoxicity and unnecessary photobleaching. These factors were particularly important when the Litmus-body complexes were sequestered inside fast-moving endosomal vesicles in living cells^51^. These properties may be valuable in tracking ADCs transiting through intracellular compartments. ADCs typically exhibit specificity for molecules on the cancer cell surface and deliver cytotoxic payload once internalized and reach acidic lysosomes. For example, drug conjugation strategies that take advantages of acid-labile linkers can release payload in low pH compartments. Traditionally, ADCs are designed to have high affinity for their target at acidic pH. More recently, “acid-switched” ADCs that instead shows high affinity for their target at neutral pH can have improved lysosomal trafficking with enhanced payload delivery and cytotoxicity^52^. In either case, Litmus-body may potentially be useful in monitoring the pH-sensitive payload delivery of ADCs, as well as to screen for ADC variants that are better trafficked into compartments of the desired pH for payload delivery^53,54^.

## ASSOCIATED CONTENT

### Supporting Information

Additional figures are available as a PDF.

## AUTHOR INFORMATION

### Author Contributions

The manuscript was written through contributions of all authors. All authors have given approval to the final version of the manuscript.

### Notes

The authors declare no competing financial interests.

## ACKNOWLEDGMENT

This investigation was supported by National Institute of Health New Innovator DP2 GM229133 (M.J.P.), National Cancer Institute U54 CA210184-01 (M.J.P. and W.R.Z.), National Institute of General Medical Sciences Ruth L. Kirschstein National Research Service Award 2T32GM008267 (M.J.C.), and National Science Foundation Graduate Research Fellowship DGE-1650441 (M.J.C.). Imaging data was acquired through the Cornell University Biotechnology Resource Center, with NYSTEM C029155 and NIH S10OD018516 funding for the shared Zeiss LSM880 microscope.

