## Supporting Information for "Litmus-Body: a Molecularly Targeted Sensor for Cell-Surface pH Measurements"

**This PDF includes:**

Supplemental Figures 1-2
Supplemental Table 1

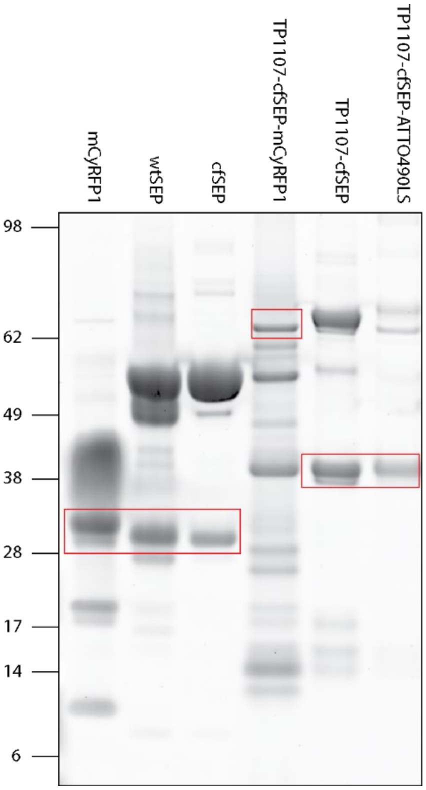


**Supplemental Figure 1.** SYPRO Ruby staining of protein elutions separated by SDS-Page. Red boxes highlight the desired protein product. Multimeric formation is observed in the solo fluorescent protein constructs.

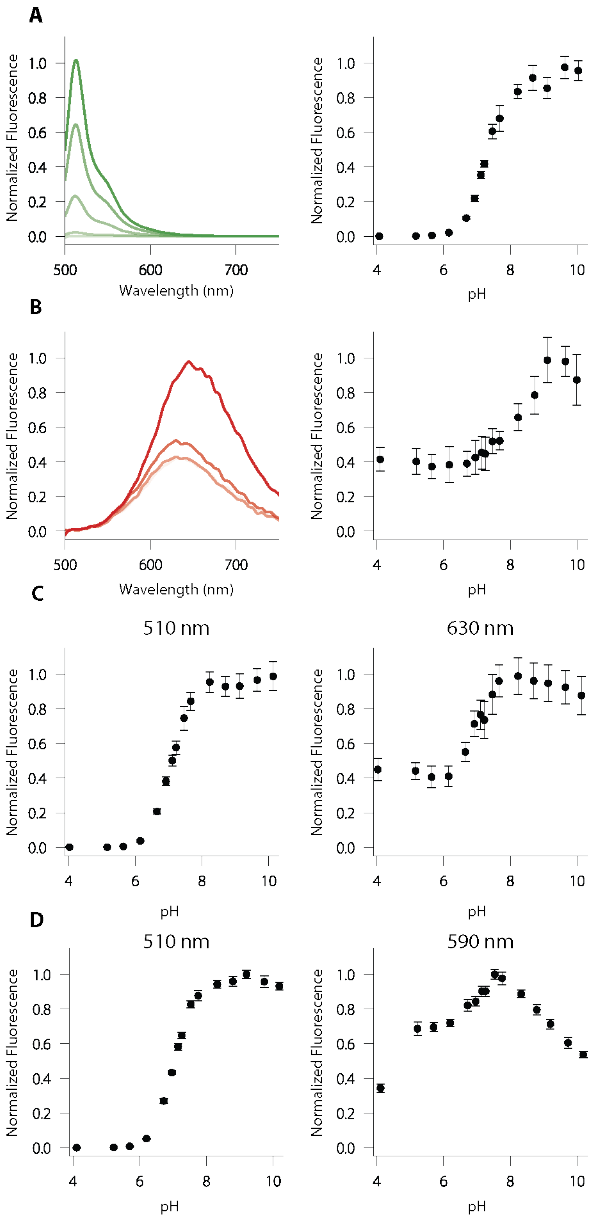


**Supplemental Figure 2.** Additional characterization of fluorophores and constructs presented. A) Spectra and pH responsiveness of wild-type SEP. B) Spectra and pH responsiveness of ATTO490LS. C) pH responsiveness at 510 nm and 630 nm respectively of the TP1107-cfSEP-ATTO490LS. D) pH responsiveness at 510 nm and 590 nm respectively of the TP1107-cfSEP-mCyRFP1.

**Supplemental Table 1.** Fit parameters of green fluorescent proteins and 510 nm emission of constructs for pH responsiveness.

| Construct | Hill Coefficient | pKa |
|---|---|---|
| SEP | 0.9 | 7.3 |
| cfSEP | 1.2 | 6.9 |
| TP1107-ATTO490LS-cfSEP | 1.4 | 7.1 |
| TP1107-mCyRFP1-cfSEP | 1.4 | 7.0 |
